# Quadratic Programming Data Descriptors for Abnormal Beat Detection in ECG Recordings

**DOI:** 10.1101/218008

**Authors:** Fayyaz ul Amir Afsar Minhas

## Abstract

This paper analyzes the efficacy of applying one class classifiers (OCCs) to the problem of abnormal beat detection in ECG. It also proposes a novel OCC called Quadratic Programming Dissimilarity representation based Data Descriptor (QPDDD). A comparison of the proposed classification technique with existing classifiers over the MIT-BIH arrhythmia database is presented. Results show that OCCs coupled with wavelet domain features present a practical, robust and scalable solution for handling inter-individual variability in ECG patterns of different types of cardiac beats. An equal error rate of 90-95% was obtained for the MIT-BIH arrhythmia database depending upon the amount of training data used. A major advantage of the proposed scheme is that it requires only normal beats during its training. Another advantage is that it is able to handle inter-individual differences in ECG morphologies as the training takes place separately for each individual.

## 1 Introduction

Discrepancies in cardiac pacemaker sites or blockages in the conduction network of the heart can lead to abnormal cardiac rhythms. An effective and extensively used non-invasive sensing modality for detecting such abnormalities is the Electrocardiogram (ECG) [1]. A system for automatically detecting abnormalities in cardiac rhythm can be extremely helpful to medical experts as it can reduce the time that an expert has to spend in analyzing the ECG. The structure of such a system is shown below.

**Figure 1.**
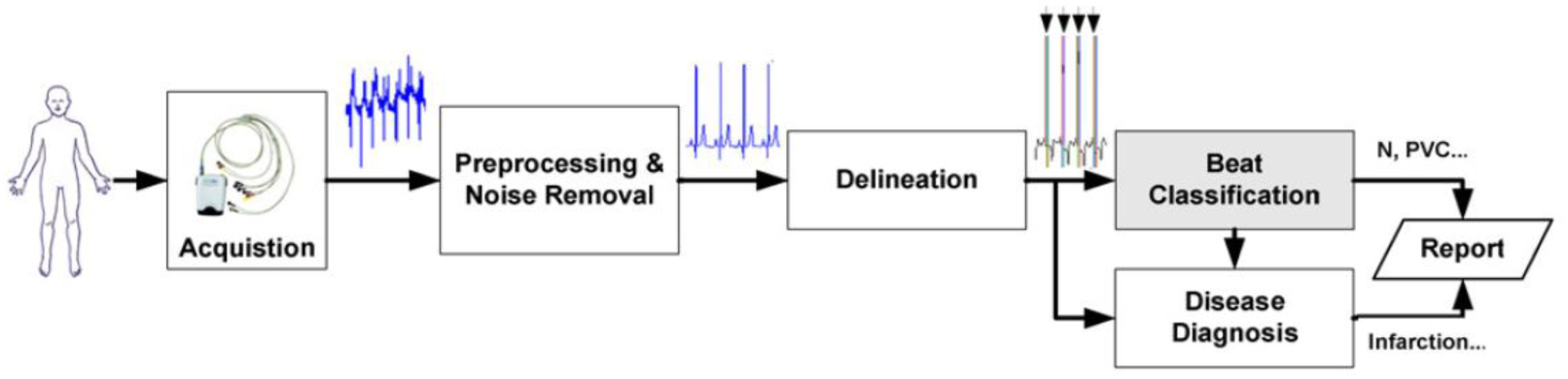
Components in a ECG Based Automated Diagnosis System

ECG is captured from a subject using acquisition hardware and is preprocessed to remove baseline variations and noise. The delineation stage is responsible for detecting and delineating different constituent parts of an ECG beat such as the P, QRS and T waves. The information extracted during the delineation phase is subsequently used in the detection of abnormal beats which is the primary focus of this paper. Effective diagnosis of complex cardiac disorders such as myocardial infarction, ischemia etc. requires that such abnormal cardiac beats be removed before parameters (such as the ST level) used in the diagnosis of these disorders are estimated. The information from an abnormal beat detector is also helpful to medical experts in identifying the type and, at times, the source of a rhythm related abnormality in the heart.

A number of methods exist in the literature for detection of abnormal beats. These include methods using features derived from Fourier Transform (FT) [2], Wavelet Transform (WT) [3], Independent Component Analysis [4], Statistical moments [5] etc. A wide variety of classification techniques such as Neural Networks [6], Support Vector Machines (SVM), Instance based classifiers [3], Neuro-Fuzzy systems [7], etc. have been employed in the literature for classifying the different types of cardiac beats.

Most of the existing research on beat classification uses both normal and abnormal beats (annotated by medical experts) in order to train a binary or multiclass classifier which can then be employed to classify a given test beat. However, ECG morphologies for the same labeled beat type across different individuals can be very different. For example, for Bundle Branch Blocks, different persons may have blockages at different locations along the bundle branch and this would cause very different patterns in the ECG for the same labeled beat type. Such differences in ECG morphologies can reduce the accuracy of conventional classifiers when the test beats come from a person whose beats were not a part of the classifier’s training set. Figure 2 illustrates this issue.

**Figure 2.**
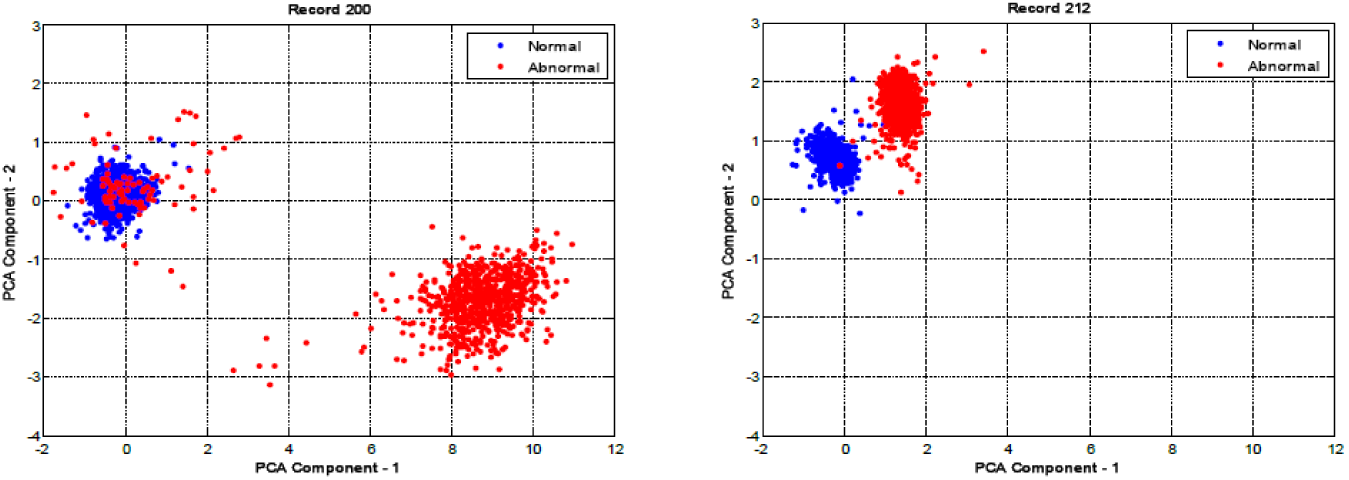
Scatter plot of principal components of time domain representation of PQRST complexes of a number of normal and abnormal beats from two different subjects (records 200 and 212) in the MIT-BIH Arrhythmia database clearly shows different data distributions for the same labeled beat type.

Such inter-individual differences in ECG beat morphologies can stem from differences in age, gender, physical condition, emotional state, genetics and family history and the exact cause of a cardiac abnormality [1]. Information regarding these peculiar differences in ECG morphologies amongst individuals has even been exploited for performing biometric identification using ECG [8] [9]. Most of the existing research in beat classification [3],[4],[5] ignores this inter-individual variability by conducting accuracy assessments of proposed schemes over individuals whose data (comprising of beats different from the ones used in testing) has already been used to train the beat classifier. However due to large intra-class (for the same labeled beat type) variability across individuals, the performance of most of existing approaches for beat classification turns out to be much lower than the reported results for data from new subjects on which the system has not been previously trained. Thus, most of the existing approaches for beat classification suffer from poor generalization across individuals.

One of the first clear mentions of the issue of high intra-class variability across individuals can be found in the paper by Peng Li et al. [10]. The method proposed in [10] trains a separate patient specific classifier based upon data from a particular individual only. It is difficult to train a conventional classifier (such as binary SVM or back-propagation neural networks etc.) for each person as a large number of both positive and negative examples would first have to be annotated by a medical expert until a training data set of substantial size is obtained for training such a classifier (as in the approach proposed in [11]). From a practical point of view, this is impossible to do as typically the number of normal beats in an ECG recording is significantly larger than abnormal beats. Such data imbalance would not only make data collection difficult but also the resulting class imbalance can impede effective learning of a conventional classifier [12]. As a consequence, Peng Li et al. have used a one-class SVM [13]. One class classifiers, such as the one-class SVM (1c-SVM), can be used to learn the concept of a target class using data from only that class for training (more details on OCC are given in section 3). In [10], a medical expert has to annotate, for each person, only a small number of normal beats which are then used to train the 1cSVM. The remaining beats are classified as normal or abnormal based upon the trained 1cSVM. This approach gives Sensitivity (Se)/Specificity (Sp) of 87.6%/95.8% over 22 subjects from the MIT-BIH arrhythmia database [14] using 38 time domain ECG features in a window around the R-peak along with the RR-interval (time difference between the R peaks of two consecutive beats). The proposed Se / Sp values are much higher than other conventional classifiers.

However, the paper by Peng Li et al. uses only a small number of subjects for evaluation. Moreover, the time domain features used in the technique are susceptible to noise. Existing research has shown that wavelet domain (or time-frequency) based features are more suitable for analyzing the inherently non-stationary ECG signal in comparison to only time domain and only frequency domain features [3]. Furthermore, one-class support vector machines for classification make a number of assumptions (discussed in section 2) about the data and the possible violations of these assumptions when dealing with real world data can reduce the accuracy of the system, Therefore, changing the type of one-class classifier can help improve the performance of the system further.

Some of the existing methods, do consider inter-individual differences in arrhythmia classification such as [15][11][16]. However, they require training data for both normal and abnormal classes during training which can limit their practical applicability in comparison to the one-class classification based approaches which are the focus of this paper.

In this paper, the focus is on different methods for one class classification and the evaluation of the efficacy of their application to the problem of automated beat classification. A new dissimilarity representation based one class classifier (called Quadratic Programming Dissimilarity representation based Data Descriptor (QPDDD)) is proposed.

## 2 One Class Classification (OCC)

This section describes the problem of OCC in detail along with some existing approaches for OCC considered in this work and the issues associated with each.

One class classifiers use training examples only from one class (called the target class) to develop a data or concept description for the target class which can then be used to determine whether a given sample belongs to the target class or not. Mathematically, the problem of OCC learning can be described as follows: given *N* training examples (*z*_*i*_,*i* = 1… *N*), find a decision function *f*(*z*;***w***) (with parameters ***w***) called the target data descriptor such that most of the target class data have *f*(*z*;***w*** ≥ 0 while most of the outliers have *f*(*z*;***w*** < 0. One-Class classification can be effective in classification problems in which one of the classes is sampled very well whereas the other class is severely under-sampled due to possible difficulties in acquisition of training examples belonging to that class. For example, while working on the design of a system for detecting faults in the operation of a machine, it may be very easy to obtain training examples for the normal behavior of the machine. On the other hand, measurement of outliers involves subjecting the machine to all possible faults that may occur which may be very expensive, if not impossible.

The problem of detecting abnormal beats in ECG is also a type of problem for which OCCs are particularly suitable since it is difficult to obtain positive examples (abnormal beats) for all possible types of abnormalities and the number of negative (or normal) training examples can usually be much larger. Moreover, as discussed earlier, there is a significant variability in beat morphologies across different individuals. This variability calls for the use of person-specific classification and usually it is very hard to collect and label training samples of abnormal beats for an individual.

One of the simplest ways to do one-class classification is to synthetically generate outlier data around the target class and then use a conventional classifier over the target class and the outlier data [17]. However, such approaches can perform poorly in high dimensional spaces. Another possible method can be to apply a probabilistic estimation method to determine the density of target class objects [18]. However such approaches require estimates for the prior probabilities and also a significantly large data set before reliable probability estimation can be done. Such problems associated with these approaches have forced the focus of research in one-class classifier design to be shifted in favor of methods that avoid estimation of complete data densities and are able to obtain the boundary of the target class directly. Such methods have minimal assumptions about the data and do not require large data sets (like probability estimation based techniques). In this paper, the focus is on non-parametric approaches that make minimal or no assumptions about the distribution of the data due to the nature of the classification problem being addressed. Some of the most popular techniques in this domain include 1cSVM [13], Support Vector Data Descriptors (SVDD) [19] and Linear Programming Data Descriptors (LPDD) [20]. A brief description of each of these methods follows.

In One-Class Support Vector Machines (1c-SVM) the decision function takes the form of a hyper-plane *f*(*z*;***w*** = ***w***^***T***^***Φ***(*z*) + *b* in a high dimensional kernel space (with the underlying implicit feature representation ***Φ***(*z*)). All training examples from the target class are constrained to lie above this hyper-plane while the weight vector ***z*** is chosen such that the hyper-plane lies at maximal perpendicular distance from the origin in an effort to maximize the margin. In the formulation of the problem the origin acts as a prior where the outlier class is assumed to lie. This assumption puts constraints on the choice of kernels that can be used in order to ensure a good closed boundary around the target class.

SVDD is closely related to 1cSVM and it attempts to find a minimum radius hyper-sphere (in a kernel space) such that target class data points are constrained to lie within that hyper-sphere. This method has been shown to be equivalent to 1c-SVM under the assumption that all training vectors have unit norm.

Another possible avenue for performing OCC is to use dissimilarity representations. The idea of using dissimilarity representations for multiclass classification problems was presented by Pękalska et al. [21]. Dissimilarity representations represent a given object only in terms of its distances/dissimilarities from other objects that belong to a representation set ***R***. This allows objects to be described directly on the basis of dissimilarities instead of features. Linear Programming Data Descriptors (LPDD) is a OCC based on dissimilarity representation of an object. The decision function used in LPDD is *f*(*z*;***w**,**R***) = ***w***^*T*^***d***(*z*_*i*_,***R*** + *b*. Here ***w*** and *b* are the linear coefficients of the hyper-plane and *d*(*z*_*i*_,***R***) = [*d*(*z*_*i*_,*r*_1_ … *d*(*z*_*i*_,*r*_|***R***|_)]^*T*^ is the vector of distances of example *z*_*i*_ from points *r*_1_…*r*_***|R|***_ ∈ ***R***. The representation set is chosen a priori and the weight vector is obtained by finding the solution to a linear constrained optimization problem in which the *L*_∞_ distance of the hyper-plane from the origin is minimized while requiring that all target class examples lie below the hyper-plane *f*(*z*;***w**,**R***) = 0. LPDD has been shown to be effective in problems even with non-metric distances which allows for their use in a wider set of problem domains than kernel based methods. One of the major problems with this approach is finding a suitable representation set **R**. Another issue is that the solution to the optimization problem in LPDD gives a sparse representation of ***w*** which may not be optimal in terms of minimizing the classification error.

A comparison of the performance of all the classifiers described above in the context of abnormal beat detection in ECG recordings is also presented in this paper.

## 3 Quadratic Programming Dissimilarity Data Description

In this section, a novel one class classifier called Quadratic Programming Dissimilarity representation based Data Descriptor (QPDDD) is presented. QPDDD, like LPDD and *k*-NNDD, is based upon dissimilarity representations of objects. However, it eliminates the difficulties in choosing the representation set which are encountered in LPDD by using only the *k*-nearest neighbors of an object in building the dissimilarity representation vector for that object. QPDDD can offer improved robustness to outliers in the training set in comparison to *k*-NNDD by solving a quadratic optimization problem that minimizes both structural and empirical risks. Henceforth the mathematical formulation of the proposed scheme is detailed.

Let 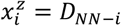 be the distance of a given example object *z* from its *i*^*th*^ nearest neighbor in the available training data. In QPDDD, an object is described on the basis of its distances 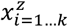 from *k* nearest training examples (excluding itself) as 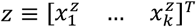. Based upon the definition of nearest neighbors, the following inequalities hold:

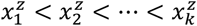

This implies that, for the *k=2* case shown in Figure 3, all training examples lie above the blue line in the dissimilarity space representation. QPDDD uses a hyper-plane ***w***^*T*^*z* − *b* = 0 (red line in Figure 3) to bound the data from above in order to obtain a data descriptor. In Figure 3, a closed boundary for the descriptor can be obtained when the hyper-plane intersects the blue line and the *x*_2_ axis. This implies that the maximum slope of the red line is +1 and the minimum slope is −∞. Thus, we have *w*_1_ + *w*_2_ ≥ 0. In general this requirement translates to the following condition.

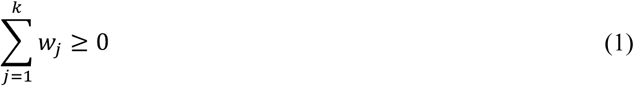

**Figure 3.**
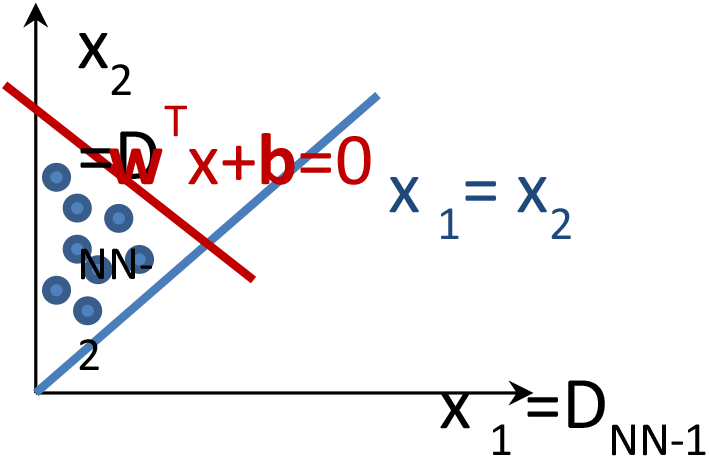
Formulation of the QPDDD in the dissimilarity space

We would like to move the line as close to the origin as possible in order to obtain a tighter boundary around the target class. The distance of the line from the origin is given by 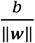. Thus the constrained optimization problem associated with the QPDDD can be written as follows.

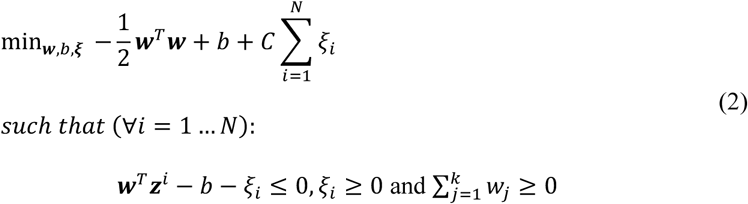

We can represent this problem using Lagrange multipliers 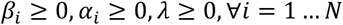 as:

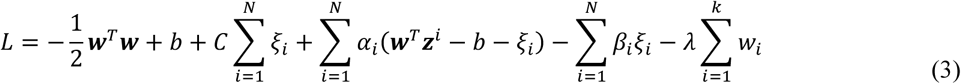

Differentiating *L* with respect to ***w***, *b* and ***ξ*** and setting the derivatives to zero we get the following dual formulation which can be solved by a standard QP solver.

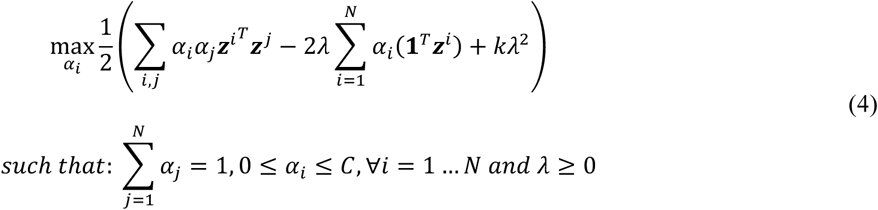

We can represent 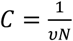 with *v* ∈ (0,1] to incorporate effects of changing the number of training examples. The parameter *v* controls the weighting of the penalty associated with constraint violation in the objective function. A large value of *v* would increase the number of margin violations leading to a tighter boundary and vice versa.

The results of applying QPDDD on some toy data sets with *k*=2 are shown below. Figure 4 shows both the original and the distance representation of a target class data set with three distinct clusters if varying sizes. It also shows the closed boundary solution obtained in the original feature space corresponding to the hyper-plane in the dissimilarity space. The distance measure used is the standard *L*_*p*_ distance with *p* = 2.

**Figure 4.**
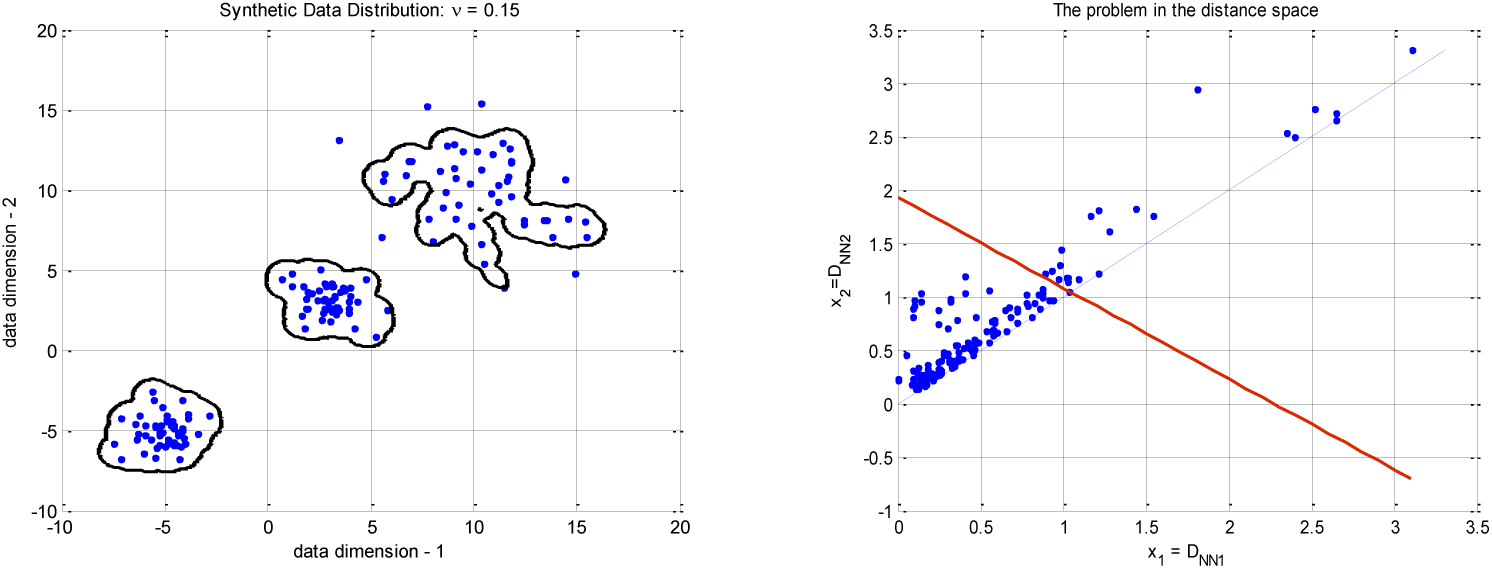
QPDDD (***p*** = **2**, ***k*** = 2, ***v*** = **0.15**) in feature (left) and dissimilarity (right) spaces

Figure 5 shows the effect of changing the number (*k*) of nearest neighbors being used in the representation over the spiral data set. As is apparent from the figure, increasing *k* increases the smoothness of the resulting boundary.

**Figure 5.**
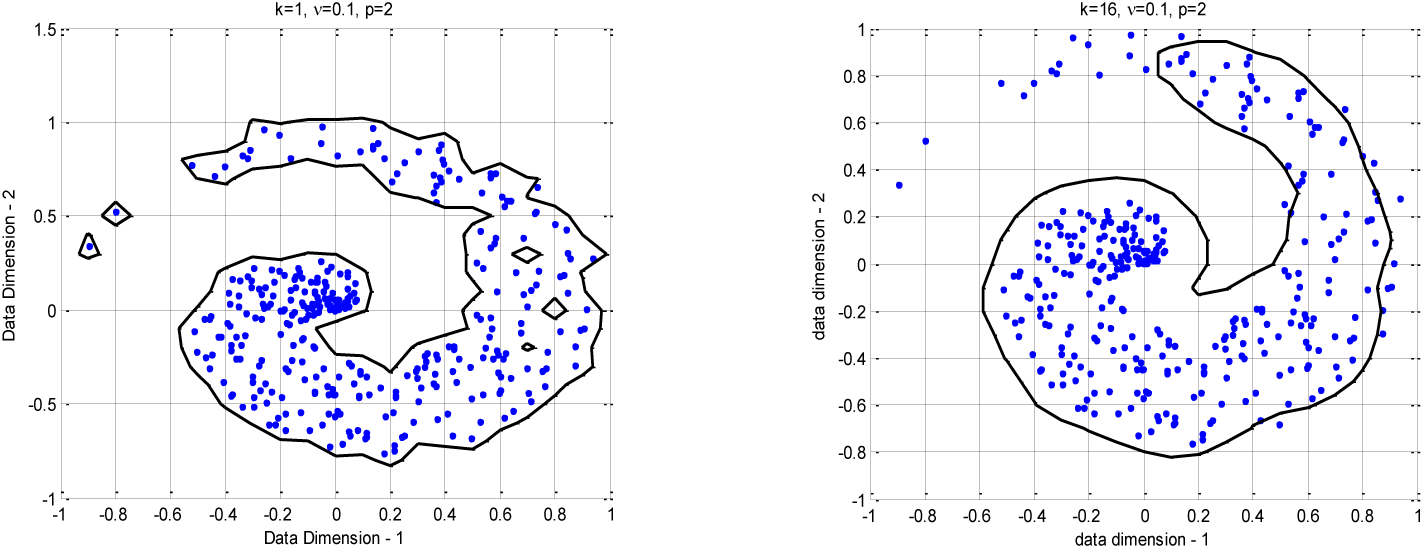
QPDDD with ***k*** = **1**(left) and ***k*** = **16** (right) with ***p*** = 2,***v*** = **0.1**.

The proposed scheme also works well with non-Euclidean (*L*_*p*=∞_) and even non-metric distances (such as *L*_*p*=0.5_) that violate the triangular inequality, as shown in Figure 6. When defining distances between objects we may come across cases in which we do not have Euclidean or even Hilbert spaces corresponding to the distance measure being used. The flexibility of QPDDD to operate over non-metric dissimilarities allows for its application to a much larger set of problem domains in comparison to kernel methods [21]. Although this increased flexibility is not utilized in the classification problem being addressed in this paper but problems such as image matching, 3D shape recognition etc. can potentially benefit from it [21].

**Figure 6.**
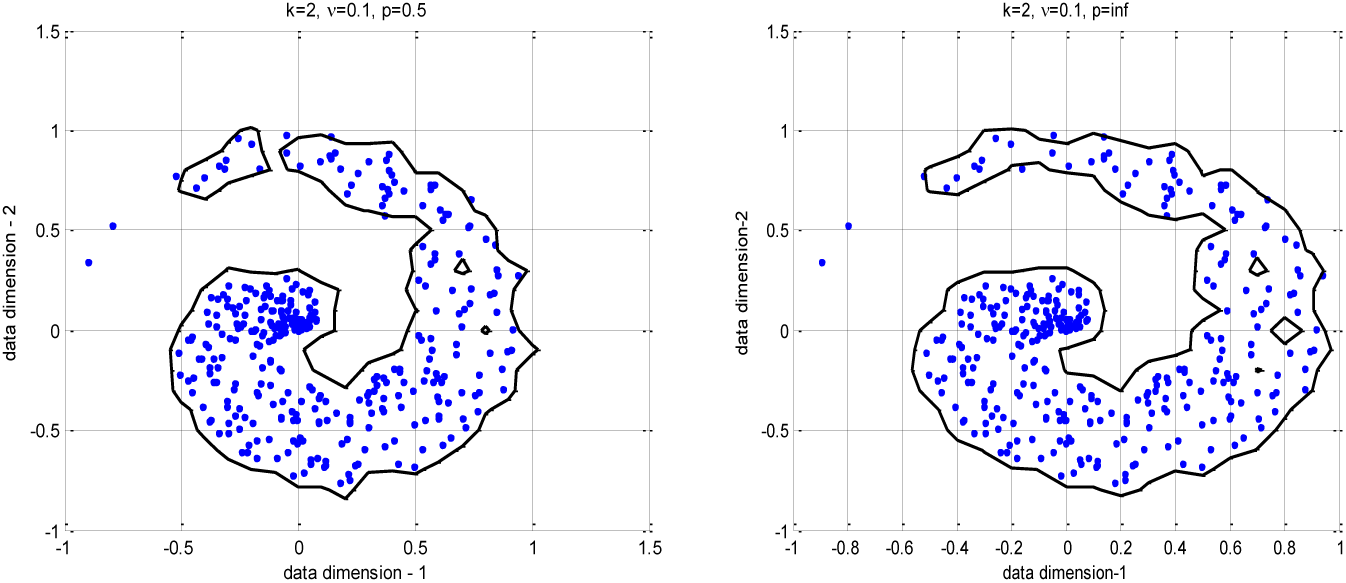
QPDDD with p=0.5 (left) and *p=*∞ (right) with ***k***=2 and ***v*** = **0.1**

## 4 Experimental Evaluation

This section compares the empirical performance of different one class classifiers discussed earlier for detection of abnormal beats in ECG recordings. The data set being used for this analysis is the MIT-BIH arrhythmia database which has become a standard in the literature for evaluating beat classification, abnormal beat detection and arrhythmia identification systems. The database contains 48 records (each of 30 minutes duration) from 47 different subjects at the Beth Israel Hospital (BIH) Arrhythmia laboratory. In this evaluation, data corresponding to ML-II channel of the ECG was used (46 out of the 48 records have data corresponding to ML-II). Four records (102, 104, 107, and 217) that include paced beats (PB) have not been used in the analysis in accordance with the guidelines of the Association for the Advancement of Medical Instrumentation (AAMI). The database consists of a total of 73,258 normal (~69.0%) and about 32,827 (~31%) abnormal beats. The dataset (henceforth called Dataset-1) comprising of the remaining 44 records is used in the analysis.

In order to compare the performance of the proposed system with the one given by Peng Li et al. [10], the proposed system has also been tested using the same data as used by Peng Li et al. Their paper uses the following records for evaluating their one-class classifier: 100, 103, 105, 113, 117, 119, 121, 123, 200, 202, 210, 212, 213, 215, 219, 221, 222, 228, 231, 232, 233, and 234. This dataset will henceforth be called Dataset-2.

In order to ensure reproducibility of the results, this section presents the details about pre-processing of the data and the procedure for feature extraction. However, we do not present a very detailed description of these processes as this is not the primary focus of this paper. The interested reader is referred to [3] for more explanation over the features and their pre-processing.

### 4.1 Pre-Processing

In the proposed scheme, signal pre-processing has been kept to a minimal (baseline removal only using a simple Finite Impulse Response (FIR) High-Pass filter with a cutoff at 0.8Hz) so that the noise-robustness of the proposed approach can be demonstrated effectively.

### 4.2 Feature Extraction

The ECG signal corresponding to each beat is divided into two segments (1 and 2) for feature extraction as shown below. Segment-1 corresponds to the signal from 0.18 seconds prior to the R-peak to the R-peak of the beat whereas Segment-2 spans the signal from the R-peak to 0.2 seconds after the R-peak. This allows a better analysis of Atrial and Ventricular components of each beat. Each segment is then decomposed using the A’trous Wavelet Decomposition and statistical features are extracted from the Approximation and Details coefficients of the decomposed signals. The complete process for extraction of these statistical features is detailed in [3]. Each beat is represented by a 21-dimensional feature vector comprising of 10 statistical features from the wavelet decomposition of each of the two segments and the RR Interval.

### 4.3 Performance Evaluation technique and Metrics

In performance analysis, each of the one class classifiers was trained with a fixed number of beats (*N*) from the start of each record and the remaining beats were used for testing. Different values of *N* were used in order to analyze the effect of training size on classifier performance. The parameters associated with different classifiers were optimized using cross validation.

The following statistical measures have been calculated to quantify the performance:

- Area under the Receiver Operating Characteristic Curve (AuROC): AuROC is closely related to the Mann-Whitney U-Test and it quantifies the number of times the classifier is able to produce a higher output for abnormal beats (positive) than normal beats (negatives). The AuROC is an excellent measure of the discrimination capability of a classifier and hence it can be a very effective measure of comparing performance of different classifiers.
- Equal Error Rate (EER): EER is the point in the ROC where the Sensitivity of the classifier becomes equal to the Specificity. It can thus be used as a practical measure of the system’s accuracy. Sensitivity is defined as 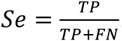. Here *TP* is the number of true positives and *FN* is the number of false negatives. Specificity is defined as 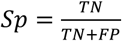. Here *TN* is the number of true negatives and *FP* is the number of false positives.

These measures provide a clearer picture in comparison to simple measures such as accuracy which are affected by the number of positive and negative examples. Moreover, providing a single specificity value at a given sensitivity is not sufficient to evaluate how a method performs in general. The measures given above were averaged for each individual in cross validation and the average values are presented.

## 5 Results and Discussion

The results of applying one class classification approaches discussed earlier to both data set 1 and 2 (D1 and D2) with different number of training beats are shown in Figure 7 in terms of AuROC and EER. Figures 7 shows that the AuROC is more than 0.90 for all the one class classifiers with EER<17% for both the training data sets with any number of training beats being considered. This shows that one class classifiers tend to be well suited to the problem under investigation.

**Fig. 7.**
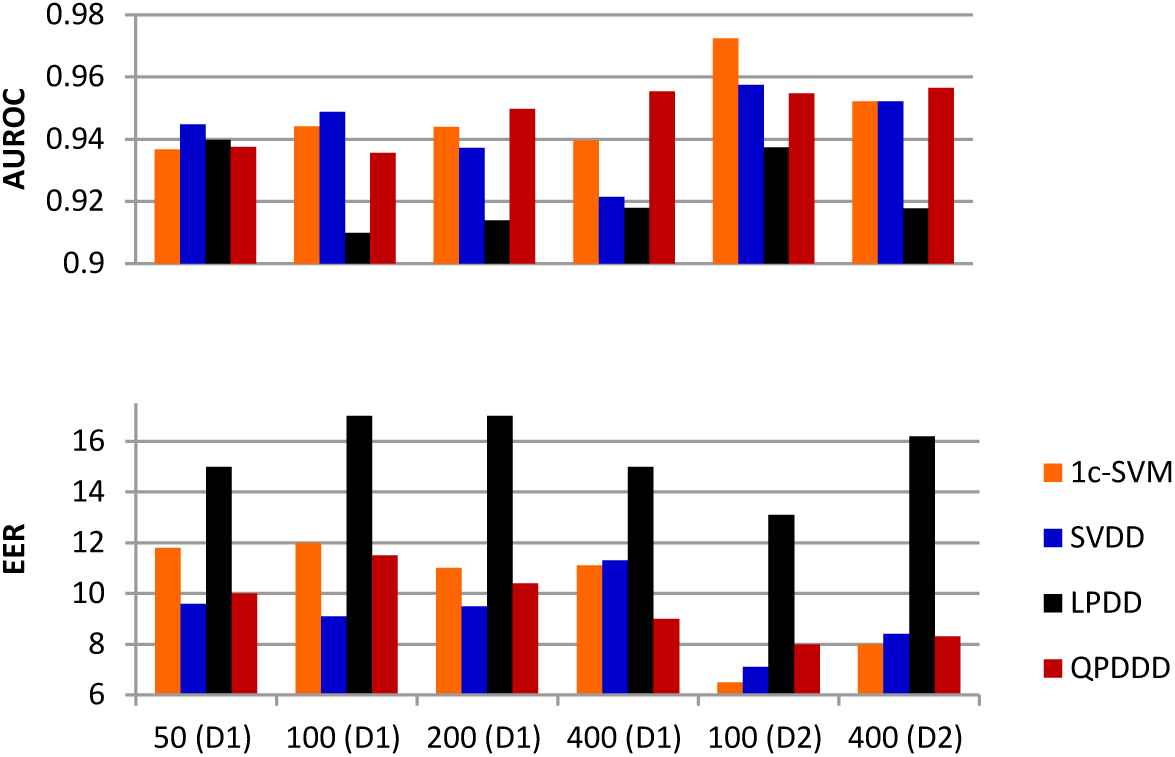
Results with first N beats used for training. Here, D1 refers to Data set 1 which consists of 46 out of the 48 records from the MIT-BIH Arrhythmia database whereas Data set 2 (D2) is composed of the same records as used in **[10]**

### 5.1 Results over Data Set 2

In comparison to the original paper by Peng Li et al., the performance of the 1cSVM (over dataset D2 with 100 training beats) with the wavelet domain features is roughly 3% better which illustrates the effectiveness of using wavelet domain features for abnormal beat detection in ECG. It must be noted that no explicit noise removal was done in this evaluation. This underlines the robustness of wavelet domain features to noise.

It must also be noted that the 1cSVM offered best results over data set 2 than all the other classifiers with N=100 training beats. However with N=400 training beats k-NNDD and QPDD performed slightly better than 1c-SVM.

### 5.2 Results over Data Set 1

Only a slight degradation in the performance of all the one class classifiers was noted when the number of subjects in the analysis was doubled when moving from D2 to D1. This clearly shows the scalability of the use of one class classifiers for the target problem.

In terms of AuROC, the performance of QPDDD is roughly the same as that of 1c-SVM and SVDD. All the classifiers show only minor changes in both EER (<2%) and AuROC (<0.02) when the amount of training data is increased from 50 to 400. This shows that one class classifiers can eliminate the need of labeling large training data sets for obtaining good performance.

### 5.3 Performance of QPDDD

Both QPDDD and LPDD are based on (different) dissimilarity representations. Amongst all the classifiers, LPDD shows significantly bad performance. Attempts were made to improve the performance of LPDD by using various different sized representation sets from the training data but no significant improvement was observed. Applying an exhaustive technique to find the optimal representation set for LPDD was impractical in our context. QPDDD, on the other hand performed significantly better than LPDD and at times even better than the kernel based methods. This clearly illustrates the effectiveness of the improved dissimilarity representation being used in QPDDD. All the reported methods show roughly similar computational efficiency.

### 5.4 Effect of using randomly selected beats for training

If the training beats are selected randomly instead of using the first *N* beats only, the performance of all the one class classifiers goes up significantly as shown Figure 8. Another observation in this regard is that in terms of EER, QPDDD now performs slightly better than all the other classifiers.

**Fig. 8.**
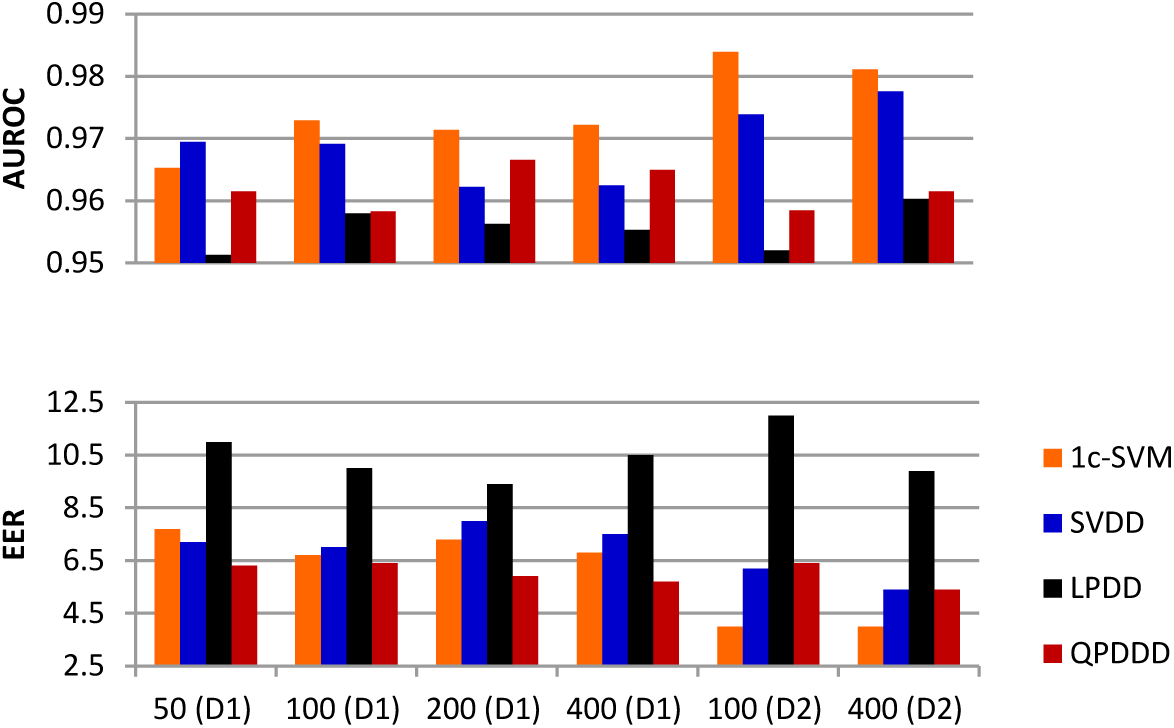
Results with randomly selected N beats for training. Here, D1 refers to Data set 1 which consists of 46 out of the 48 records from the MIT-BIH Arrhythmia database whereas Data set 2 (D2) is composed of the same records as used in **[10]**

## 6 Conclusions and Future Work

We have seen that one class classifiers present a very viable and practical solution for handling the inter-individual differences in ECG beat morphologies for abnormal beat detection. It has also been shown that the novel QPDDD proposed in the paper offers excellent performance in comparison to other methods. QPDDD has also shown significantly better results than existing dissimilarity representation based LPDD. Future work will focus on using a direct dissimilarity representation that would allow us to use QPDDD over raw ECG signals without feature extraction. Currently, the proposed scheme only discriminates between normal and abnormal beats due its reliance on One Class Classification. In future, a more detailed classification of different types of arrhythmias can be considered through a cascaded classification scheme in which the abnormal beats are first detected using QPDDD and then a conventional multi-class classifier is used to classify those abnormalities into specific categories.

